# Posterior Staphyloma-associated vitreous proteomic remodeling in high myopia

**DOI:** 10.64898/2026.11.16.767054

**Authors:** Kimberley Delaunay, Clément Guillou, Emilie Picard, Laurette Fernandez, Laurent Jonnet, José Ruiz Moreno, Jorge Ruiz-Medrano, Pascal Cosette, Olivier Cases, Francine Behar Cohen

## Abstract

Posterior staphyloma is a hallmark of pathologic myopia, but the cellular and molecular changes of the overlying retina remain poorly understood. We combined the histopathology of human eyes with posterior staphyloma with label-free proteomics of vitreous from non-myopic eyes (n = 4) and highly myopic eyes without (n = 3) or with posterior staphyloma (n = 7) and measured soluble vitreous iron in an independent series of 31 eyes. In the staphyloma, the retina collapsed at the transition zones into a vimentin- and GFAP-positive glial scar containing Iba1-positive microglia, with iron deposits, increased ferritin and disorganized scleral collagen. The vitreous proteome was not globally altered, but protein modules defined a priori to mirror these lesions revealed an activation of the classical complement pathway and an increased ferritin-to-transferrin ratio, both specific to staphylomatous eyes, whereas neuron- and photoreceptor-derived proteins were already decreased in highly myopic eyes without staphyloma. Soluble vitreous iron was lower in staphylomatous eyes than in non-myopic eyes and in highly myopic eyes without staphyloma, independently of total protein content. These findings suggest that complement activation and iron sequestration characterize the retinal degeneration associated with posterior staphyloma, and that the vitreous provides a window on these processes.

## Introduction

Myopia is the most common refractive error worldwide and its prevalence is rising rapidly. By 2050, an estimated 4.8 billion people (49.8% of the world population) will be myopic and 938 million (9.8%) highly myopic^1^. High myopia can progress to pathologic myopia, defined by myopic maculopathy and/or posterior staphyloma, which has become a leading cause of irreversible visual impairment^2^.

A posterior staphyloma is an outpouching of the posterior wall of the eye, with a radius of curvature smaller than that of the surrounding eye wall and is considered a hallmark of pathologic myopia^2^. Curtin first classified posterior staphylomas into ten types on ophthalmoscopic grounds^3^; Ohno-Matsui later proposed a simplified classification based on three-dimensional magnetic resonance imaging and wide-field fundus imaging^4^. Beyond its anatomical characteristics, posterior staphyloma is strongly associated with the development and severity of myopic maculopathy. Ruiz-Moreno et al. (2023) highlighted its importance as a determining factor in myopic maculopathy, emphasizing that posterior ocular deformation is clinically relevant beyond axial elongation alone.

Histologically, myopic axial elongation is associated with thinning of the sclera and choroid, most marked at the posterior pole, and with defects of Bruch’s membrane in the macular and parapapillary regions^5^. In enucleated human eyes, posterior staphylomas are characterized by marked scleral thinning and spatially correlated Bruch’s membrane defects, whereas retinal pigment epithelium density, Bruch’s membrane thickness and choriocapillaris thickness do not differ from those of highly myopic eyes without staphyloma^6^. These descriptions have focused on the eye wall; the cellular response of the overlying neural retina, and in particular of its glial and immune cells, remains poorly characterized.

Several lines of evidence point to an inflammatory and degenerative component in pathologic myopia. Aqueous humour levels of complement proteins of the classical and alternative pathways are higher in highly myopic eyes, are highest in eyes with posterior staphyloma and chorioretinal atrophy, and correlate with thinning of the macular retina^7^. Complement factor H is also elevated in the aqueous humour of highly myopic eyes, particularly those with choroidal atrophy^8^. In the retina, the initiator of the classical pathway, C1q, drives microglial activation and neurodegeneration^9^. Iron is another candidate mediator of retinal damage: its homeostasis is tightly regulated locally in the retina, and excess iron catalyses the formation of reactive oxygen species and has been implicated in several retinal degenerations^10^·^11^. Whether iron accumulates in the staphylomatous retina is unknown.

The vitreous humour lies in direct contact with the inner retina and is sampled routinely during vitrectomy, making it an accessible proxy of the retinal microenvironment. Label-free proteomic studies have compared the vitreous of eyes with pathologic myopia with that of non-myopic eyes, reporting changes in antioxidant proteins^12^ and in proteins related to chorioretinal cell apoptosis and scleral extracellular matrix synthesis^13^, and a reanalysis of published myopia proteomic datasets highlighted innate immunity and inflammation^14^. However, these studies did not distinguish eyes with posterior staphyloma from highly myopic eyes without staphyloma, and none related vitreous changes to the histopathology of the staphylomatous retina.

Here, we combined the histopathological analysis of human eyes with posterior staphyloma with label-free proteomic profiling of vitreous from non-myopic eyes and from highly myopic eyes with or without staphyloma. We asked whether the staphyloma is associated with a distinct vitreous signature, and whether this signature reflects the neuronal, glial, microglial and iron-related alterations observed in the staphylomatous retina.

## Results

### Histopathology of the posterior staphyloma

Serial Nissl-stained sections cut along the temporal–nasal axis of highly myopic eyes [authors: number of eyes, origin] delineated, within the same eye, the temporal retina, a posterior staphyloma, the macula, a peripapillary staphyloma and the optic nerve head, separated by transition zones (Supplementary Fig. S1a). At the temporal transition zone, the laminated retina collapsed abruptly: the retinal pigment epithelium (RPE) ended, and the retina separated from the underlying choroid (Fig. 1a, double-headed arrow). Within the posterior staphyloma, the neural retina was reduced to a thin tissue containing a single, discontinuous row of nuclei, lying on an attenuated, pigment-laden choroid, with no identifiable photoreceptor layer or RPE (Fig. 1b,c). In contrast, the macula, located between the posterior and peripapillary staphylomas, retained its laminar organization, although the neural retina was detached from the RPE (Fig. 1d).

**Figure 1.**
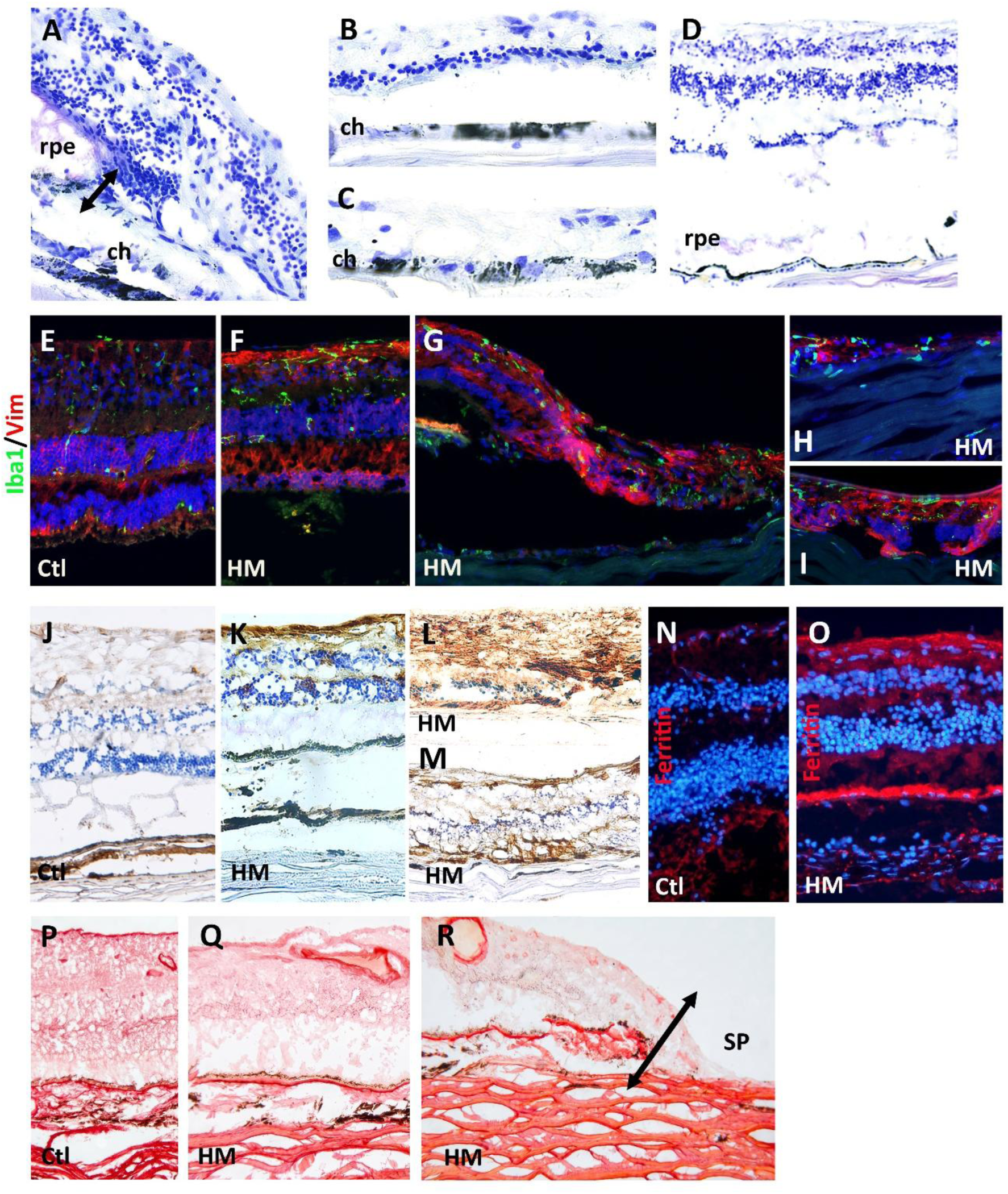
Histopathology of the posterior staphyloma. (**a**–**d**) Nissl staining. (**a**) Temporal transition zone: abrupt collapse of the laminated retina; the double-headed arrow marks the separation between the retina and the choroid where the RPE ends. (**b**,**c**) Posterior staphyloma: residual retina reduced to a discontinuous row of nuclei over a thin, pigmented choroid. (**d**) Macula between the posterior and peripapillary staphylomas: preserved lamination with detachment of the neural retina from the RPE. (**e**–**i**) Iba1 (green) and vimentin (red) immunofluorescence, nuclei in blue: control retina (**e**), retina of a highly myopic eye (**f**), transition zone (**g**) and staphyloma (**h**,**i**). (**j**–**m**) Perls staining in a control retina (**j**) and in highly myopic eyes with posterior staphyloma (**k**–**m**) [authors: specify the region of panels k–m]. (**n**,**o**) Ferritin immunofluorescence (red), nuclei in blue, in a control (**n**) and a highly myopic eye with posterior staphyloma (**o**). (**p**–**r**) Sirius red staining of collagen in a control eye (**p**) and in highly myopic eyes (**q**,**r**); (**r**) edge of the posterior staphyloma, the double-headed arrow marking [authors to specify]. ch, choroid; rpe, retinal pigment epithelium; SP, staphyloma; Ctl, control; HM, high myopia. Scale bars: [authors to add].

In control retina, vimentin labelled the radial processes of Müller cells, and Iba1-positive microglia were sparse and ramified (Fig. 1e). In highly myopic eyes, vimentin immunoreactivity increased, particularly in the inner retina and the outer plexiform layer, and Iba1-positive cells were more numerous in the inner retina (Fig. 1f). At the transition zone, the collapsed retina formed a dense vimentin-positive tissue containing numerous Iba1-positive cells (Fig. 1g; Supplementary Fig. S1b). In the staphyloma, the residual tissue lining the sclera was vimentin-and GFAP-positive and contained Iba1-positive cells (Fig. 1h,i; Supplementary Fig. S1c), consistent with a glial scar.

In control retina, Perls staining was faint in the neural retina and found mainly in the choroid (Fig. 1j). In highly myopic eyes with posterior staphyloma, iron deposits were present within the neural retina, at its inner surface and in the retinal layers (Fig. 1k,m), and were dense in the collapsed staphylomatous tissue (Fig. 1l); along the temporal–nasal axis, they extended from the posterior staphyloma to the peripapillary staphyloma and were most marked around the optic nerve head (Supplementary Fig. S1d). Accordingly, ferritin immunoreactivity, weak in the neural retina of control eyes and mainly choroidal (Fig. 1n), was strongly increased in highly myopic eyes, at the inner retinal surface and as a continuous band in the outer retina (Fig. 1o). Sirius red staining showed compact, densely packed collagen lamellae in the sclera of control eyes (Fig. 1p). In highly myopic eyes, the scleral lamellae were thinner and separated by wide interlamellar spaces (Fig. 1q), a disorganization most marked beneath the staphyloma, where the choroid was attenuated and the residual retina lay on loosened scleral collagen (Fig. 1r, double-headed arrow). Together, these observations show that the posterior staphyloma combines neuronal loss, glial scarring, microglial accumulation, iron overload and disorganization of the scleral collagen, which we next sought to detect in the vitreous.

### Study population

Vitreous samples from 17 eyes were submitted to label-free LC-MS/MS: 5 non-myopic controls, 5 highly myopic eyes without documented staphyloma (MS−) and 7 highly myopic eyes with posterior staphyloma (MS+). Following the quality-control criteria of the proteomics facility, one control (low intra-group reproducibility) and two MS− samples (low alignment score and high normalization factor, respectively) were excluded, leaving 4 controls, 3 MS− and 7 MS+ eyes for analysis (Table 1; Supplementary Table S1). Controls underwent vitrectomy for intraocular lens subluxation (n = 2) or epiretinal membrane (n = 2). Age (median 62, 68 and 67 years, respectively) and sex distribution (3/4, 3/3 and 5/7 women) were comparable across groups. All MS+ eyes had a posterior staphyloma; the staphyloma type was documented in four eyes (types 1, 3, 7 and 9 following Curtin classification).

**Table 1.** Characteristics of the eyes analysed by vitreous proteomics.

| Characteristic | Controls (n = 4) | High myopia without staphyloma, MS− (n = 3) | High myopia with staphyloma, MS+ (n = 7) |
|---|---|---|---|
| Age, years, median (range) | 62 (56–68) | 68 (66–77) | 67 (45–77) |
| Women, n | 3 | 3 | 5 |
| Axial length, mm, median (range) | 22.8 (21.8–23.8), n = 2 | 25.35, n = 1 (−10 D) | 29.4 (26.7–30.8), n = 4 |
| Posterior staphyloma | None | None documented | 7/7 (types 1, 3, 7, 9; 3 not typed) |
| Indication for vitrectomy | IOL subluxation (2), epiretinal membrane (2) | Epiretinal membrane (3) | Myopic traction maculopathy/foveoschisis (3), IOL subluxation (1), IOL exchange (1), vitreomacular traction (1), macular hole (1) |

IOL, intraocular lens. Excluded by the proteomics facility before analysis: one control and two MS− samples (see Supplementary Table S1).

### The vitreous proteome of eyes with posterior staphyloma

Across all samples, 471 proteins were identified at a false discovery rate below 1%. After exclusion of single-peptide identifications, 345 proteins were retained for quantitative comparison of control and MS+ vitreous (Supplementary Data 1). Principal component analysis did not separate the two groups (PC1, 30% of variance; PC2, 17%; Fig. 2a), indicating that the staphyloma does not globally remodel the vitreous proteome.

**Figure 2.**
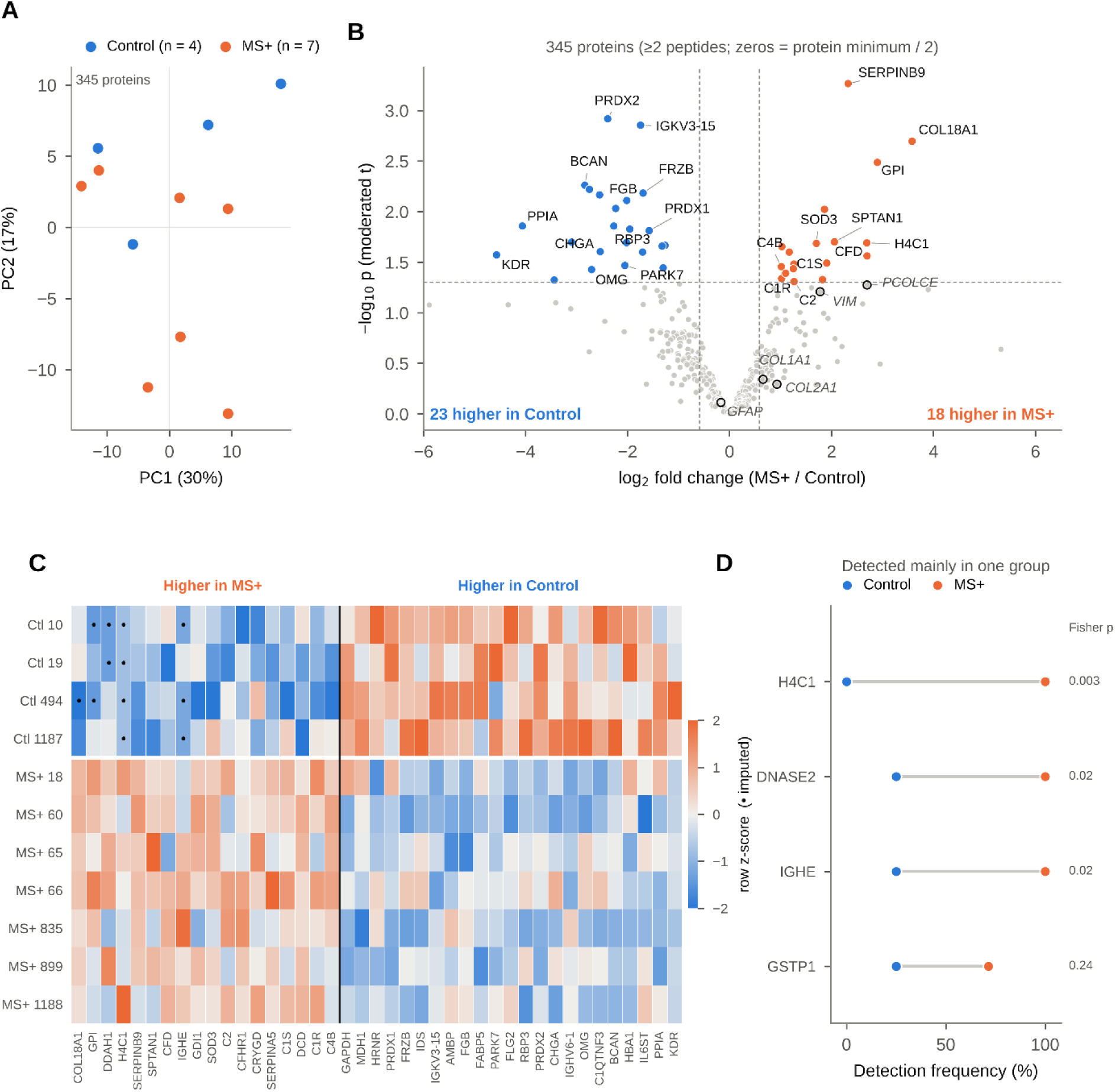
The vitreous proteome in pathological myopia with posterior staphyloma. (**a**) Principal component analysis of the 345 proteins identified by at least two peptides; controls (blue, n = 4) and MS+ eyes (orange, n = 7). (**b**) Volcano plot (moderated t-test). Dashed lines, p = 0.05 and fold change = 1.5; no protein reached a Benjamini–Hochberg false discovery rate of 5%. Open circles mark glial and fibrillar collagen markers matching the histological stains. (**c**) Heatmap of the 41 regulated proteins (row z-scores of log2 abundance), ordered by fold change; dots mark zero values replaced by half the protein minimum. (**d**) Proteins detected mainly in one group; points give the detection frequency per group; Fisher’s exact test.

Forty-one proteins nevertheless differed between groups (moderated t-test p < 0.05, fold change ≥ 1.5), 18 being more abundant and 23 less abundant in MS+ vitreous (Fig. 2b,c). None reached a Benjamini–Hochberg false discovery rate of 5% (lowest q = 0.16). The largest increases concerned serpin B9 (5.0-fold, p < 0.001), collagen XVIII (p = 0.002) and glucose-6-phosphate isomerase (p = 0.003). The largest decreases concerned peroxiredoxin-2 (0.19-fold, p = 0.001), brevican (0.14-fold, p = 0.005), frizzled-related protein 3 (0.31-fold, p = 0.007), fibrinogen β chain (0.25-fold, p = 0.008) and interphotoreceptor retinoid-binding protein (RBP3; 0.21-fold, p = 0.014). Three proteins were detected almost exclusively in MS+ vitreous: histone H4 (7/7 MS+ versus 0/4 control eyes, Fisher’s exact test p = 0.003), deoxyribonuclease II and immunoglobulin ε heavy chain (7/7 versus 1/4, p = 0.024 each; Fig. 2d).

### A vitreous signature that mirrors retinal histopathology

To relate vitreous changes to the tissue alterations shown in Fig. 1, we scored nine protein modules defined a priori by cell type or function, each matched where possible to a histological stain (Fig. 3a; Supplementary Data 1). Every module member identified by at least two peptides was included, regardless of its individual result.

**Figure 3.**
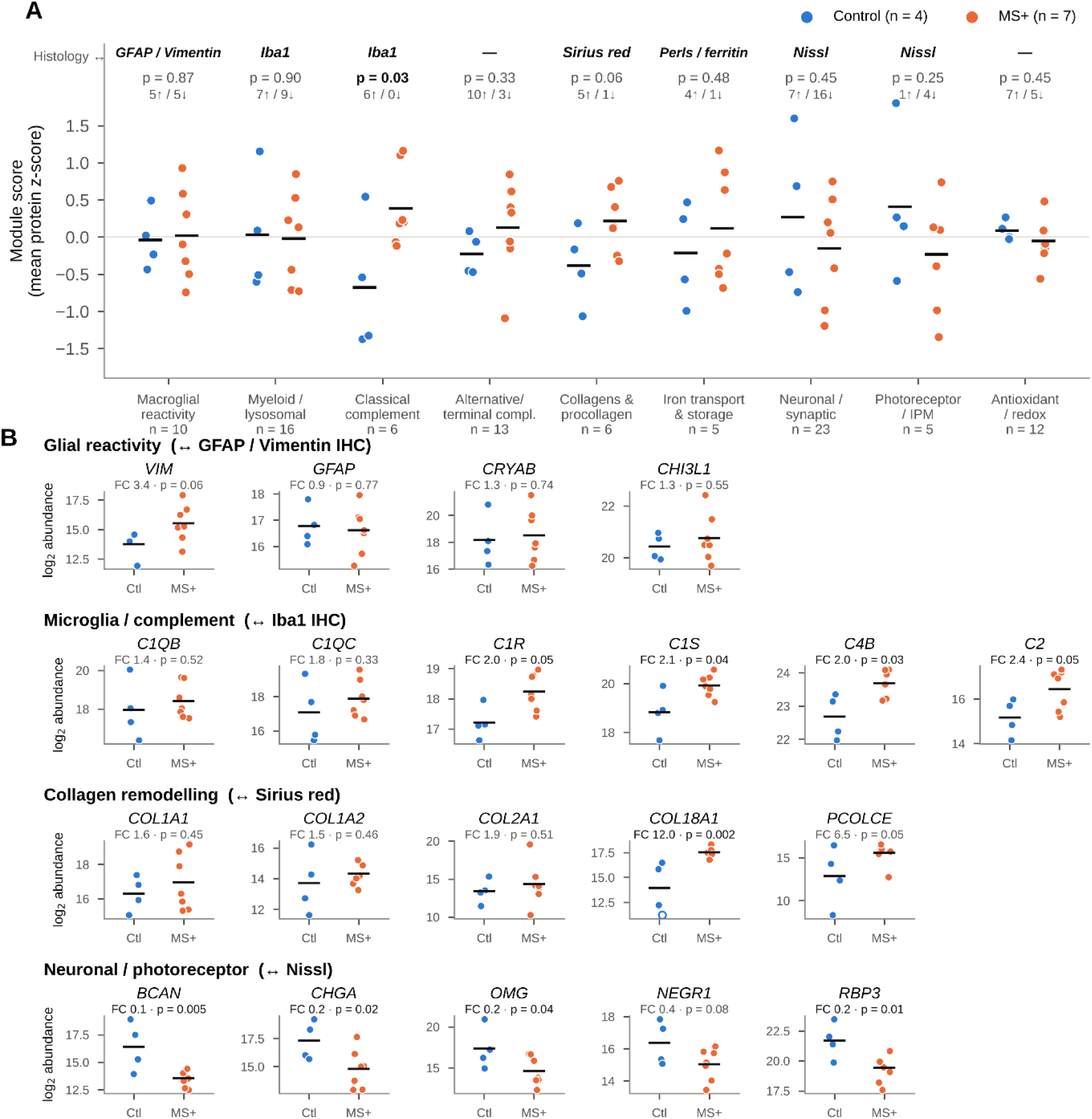
A vitreous signature aligned with retinal histopathology. (**a**) Scores of nine a-priori protein modules (mean protein z-score per eye; bars, group means). Above each module: the corresponding histological stain, exact permutation p value and number of members increased or decreased in MS+ eyes. (**b**) Marker proteins arranged as in Fig. 1: glial reactivity (GFAP/vimentin), microglia and complement (Iba1), collagen remodelling (Sirius red) and neuronal/photoreceptor proteins (Nissl). Points, individual eyes (log2 normalized abundance; open circles, replaced zero values); bars, means; fold change (MS+/control, geometric means) and moderated t-test p value above each panel.

The classical complement module was consistently increased in MS+ vitreous (6/6 members increased; module score difference +1.06, exact permutation p = 0.033). This was driven by C4B (2.0-fold, p = 0.03), C1s (2.1-fold, p = 0.04), C1r (2.0-fold, p = 0.05) and C2 (2.4-fold, p = 0.05), whereas C1q B and C chains increased non-significantly (Fig. 3b). As C1q is produced mainly by retinal microglia and initiates this cascade, this change parallels the accumulation of Iba1-positive microglia observed in the staphylomatous retina (Fig. 1e–i). The collagen module tended to increase (5/6 members increased; +0.60, p = 0.064), with collagen XVIII (p = 0.002) and procollagen C-endopeptidase enhancer 1 (6.5-fold, p = 0.05), in keeping with the disorganization of scleral collagen revealed by Sirius red staining (Fig. 1p–r).

Conversely, neuron-derived proteins tended to decrease (16/23 module members decreased), including brevican (0.14-fold, p = 0.005), chromogranin A (0.17-fold, p = 0.02) and oligodendrocyte-myelin glycoprotein (0.15-fold, p = 0.04), together with the photoreceptor-derived RBP3 (0.21-fold, p = 0.014). These changes are consistent with the neuronal loss seen on Nissl staining, although neither the neuronal (p = 0.45) nor the photoreceptor module (p = 0.25) differed significantly. Among glial markers, vimentin rose 3.4-fold (p = 0.06) whereas GFAP was unchanged (p = 0.77), and the macroglial and myeloid/lysosomal modules did not differ between groups. No module remained significant after correction across the nine modules (q ≥ 0.29); these findings are therefore hypothesis-generating.

### Changes specific to the staphyloma versus shared with high myopia

To distinguish the effect of the staphyloma from that of axial elongation, we included the three highly myopic eyes without documented staphyloma (MS−), quantified in the same run and with identical normalization (Supplementary Fig. S3). The decrease in neuron-and photoreceptor-derived proteins was already present in MS− vitreous (brevican 0.20-fold, p = 0.04; oligodendrocyte-myelin glycoprotein 0.09-fold, p = 0.04 versus controls) and did not deepen further in MS+ eyes, suggesting that it is associated with high myopia itself. Collagen XVIII and procollagen C-endopeptidase enhancer 1 followed the same pattern. By contrast, complement factor D (12-fold, p < 0.001), C1r (2.5-fold, p = 0.02), ferritin heavy chain (15-fold, p = 0.02) and serpin B9 (2.5-fold, p = 0.04) were higher in MS+ than in MS− vitreous, and the classical complement score followed a control < MS− < MS+ gradient (MS− versus MS+, p = 0.29). Thus, the complement and iron-storage changes that mirror Iba1 and Perls staining appear specific to the staphyloma, whereas vitreous markers of neuronal loss are shared with high myopia. Given the size of the MS− group, these comparisons are exploratory.

### Iron-handling proteins and corticosteroid pathway

Iron accumulation, shown by Perls staining and increased ferritin immunoreactivity in the staphylomatous retina (Fig. 1j–o), had a counterpart in the vitreous proteome (Fig. 4d). Ferritin heavy chain was 15-fold higher in MS+ than in MS− vitreous (p = 0.02) and tended to be higher than in controls (6.1-fold, p = 0.06), whereas transferrin tended to decrease (0.7-fold, p = 0.25). The ferritin-to-transferrin ratio was consequently higher in MS+ vitreous than in both control (Mann–Whitney p = 0.024) and MS− vitreous (p = 0.017). Vitreous ferritin heavy and light chains correlated with the classical complement score (Spearman ρ = 0.65 and 0.63, p = 0.03 and 0.04, n = 11), consistent with iron sequestration by activated microglia. Haemoglobin α chain (0.1-fold, p = 0.02) and α1-microglobulin (0.26-fold, p = 0.015) were lower in MS+ vitreous, while haptoglobin and haemopexin were unchanged (Supplementary Fig. S2).

**Figure 4.**
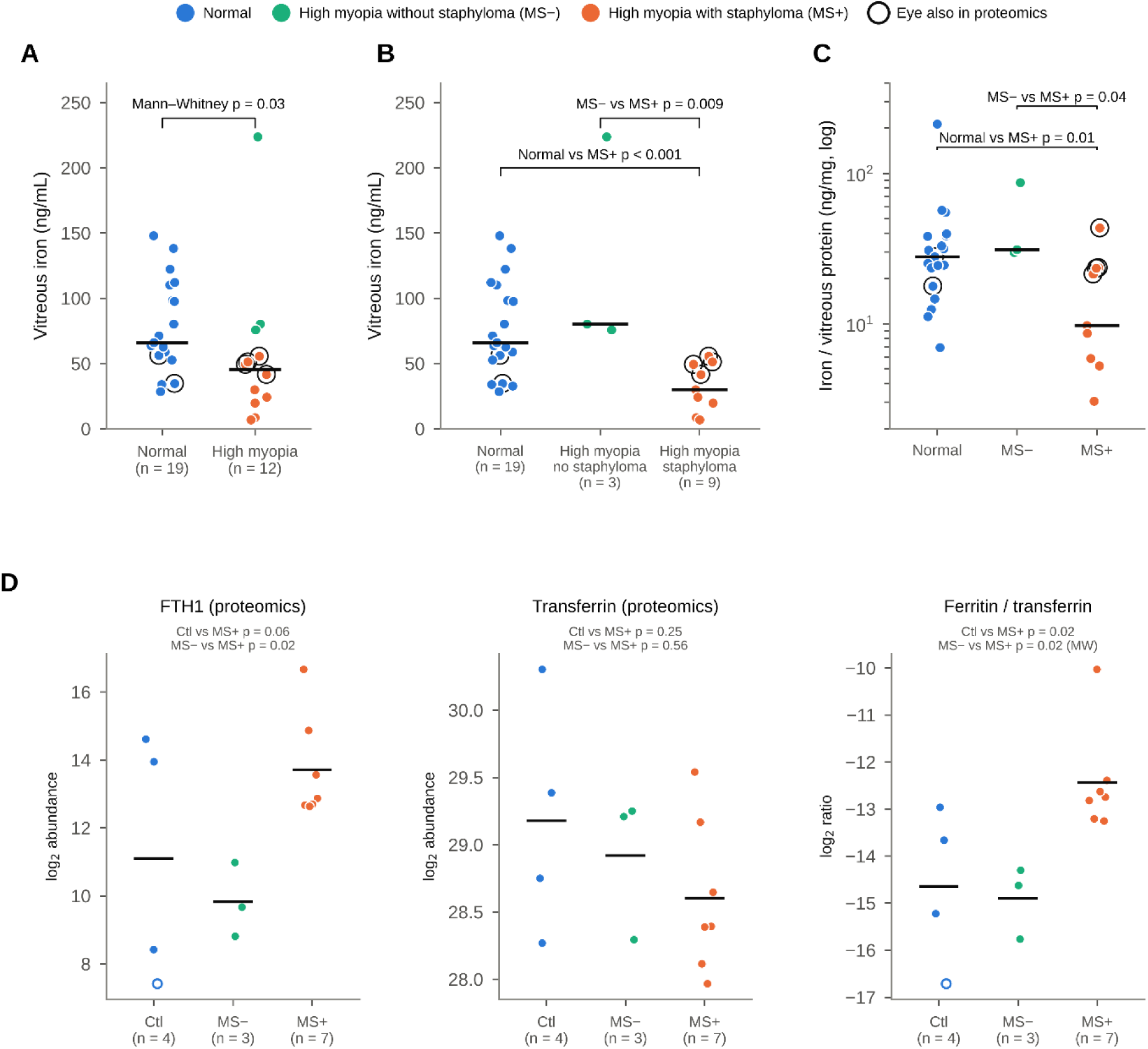
Iron handling in the myopic eye. (**a**–**c**) Soluble vitreous iron measured in a single assay batch. (a) Non-myopic (n = 19) versus highly myopic eyes (axial length ≥ 26 mm or posterior staphyloma; n = 12). (b) Same data according to staphyloma status: high myopia without (MS−, n = 3) or with posterior staphyloma (MS+, n = 9). (c) Iron normalized to total vitreous protein (log scale). Circled points, eyes also included in the proteomic analysis; bars, medians; Mann–Whitney tests. (**d**) Ferritin heavy chain, transferrin and log2 ferritin-to-transferrin ratio in the proteomic series (controls n = 4, MS− n = 3, MS+ n = 7). Bars, means; moderated t-test from a three-group model, or Mann–Whitney test for the ratio; open circles, replaced zero values.

In an independent series of 31 eyes assayed in a single batch (19 non-myopic and 12 highly myopic eyes; Supplementary Table S1), soluble vitreous iron was lower in highly myopic than in non-myopic eyes (median 45 versus 66 ng/mL, Mann–Whitney p = 0.027; Fig. 4a). This decrease was confined to eyes with posterior staphyloma (n = 9; median 30 ng/mL, interquartile range 20–49), which differed from both non-myopic eyes (p < 0.001) and highly myopic eyes without staphyloma (n = 3; median 80 ng/mL; p = 0.009), whereas highly myopic eyes without staphyloma did not differ from non-myopic eyes (p = 0.27; Fig. 4b). Excluding one MS− eye with an outlying value (224 ng/mL) did not change this result (p = 0.036). Total vitreous protein concentration did not differ between groups (Kruskal–Wallis p = 0.58), and the decrease persisted after normalization of iron to protein content (median 9.7 ng/mg in MS+ eyes versus 27.9 ng/mg in non-myopic eyes, p = 0.012, and 31.1 ng/mg in MS− eyes, p = 0.036; Fig. 4c). Among highly myopic eyes, vitreous iron was not significantly correlated with axial length (Spearman ρ = −0.37, p = 0.24). Staphylomatous eyes thus combined low soluble vitreous iron with increased vitreous ferritin and tissue iron accumulation.

In contrast, the corticosteroid pathway showed no change. Corticosteroid-binding globulin was stable (1.1-fold, p = 0.81), and glucocorticoid-responsive proteins (α1-acid glycoproteins 1 and 2, α1-antichymotrypsin, zinc-α2-glycoprotein, apolipoprotein D and angiotensinogen) did not differ as a group (p = 0.57; Supplementary Fig. S2).

## Discussion

This study combines, for the first time to our knowledge, the histopathology of the staphylomatous retina with the vitreous proteome of eyes with and without posterior staphyloma. In the tissue, the staphyloma was characterized by an abrupt collapse of the retina at the transition zones, a residual retina reduced to a vimentin- and GFAP-positive glial scar, an accumulation of Iba1-positive microglia an iron overload revealed by Perls staining and ferritin immunoreactivity, and a disorganization of the scleral collagen lamellae on Sirius red staining. The vitreous did not show a global proteomic shift, and no single protein survived correction for multiple testing; but modules defined a priori to mirror the histological stains revealed a coherent signature: activation of the classical complement pathway, an increased ferritin-to-transferrin ratio and a decrease in neuron- and photoreceptor-derived proteins. The comparison with highly myopic eyes without documented staphyloma suggests that the complement and iron changes are specific to the staphyloma, whereas the loss of neuronal proteins is shared with high myopia.

The most consistent vitreous change was the increase of the classical complement cascade, with all six quantified members, from C1q to C2, increased in staphylomatous eyes. C1q, the initiator of this pathway, is a primary mediator of microglial activation and neurodegeneration in the injured retina^9^, and its downstream components rose in parallel with the microglial accumulation observed histologically. Our findings extend aqueous humour studies showing that classical and alternative complement proteins are elevated in highly myopic eyes, highest in eyes with posterior staphyloma and chorioretinal atrophy, and associated with thinning of the macular retina^7^, and that complement factor H is increased in highly myopic eyes with choroidal atrophy^8^. Compared with highly myopic eyes without staphyloma, complement factor D was 12-fold higher in staphylomatous vitreous, whereas factors H and I tended to be lower, suggesting an imbalance between amplification and regulation of the alternative pathway that deserves confirmation in a larger series.

The second axis concerns iron. The staphylomatous retina accumulated Perls-positive iron and ferritin, and vitreous ferritin increased relative to transferrin. Iron catalyses the formation of reactive oxygen species and excess retinal iron has been implicated in several retinal degenerations^10^·^11^. In mice lacking the ferroxidases ceruloplasmin and hephaestin, retinal iron accumulation is accompanied by oxidative stress and complement activation and leads to retinal degeneration^15^, and post-mortem maculae with age-related macular degeneration contain more iron than age-matched controls^16^. The correlation we observed between vitreous ferritin and the classical complement score, together with the presence of Iba1-positive cells in iron-laden regions, raises the hypothesis that activated microglia sequester iron in the staphylomatous retina. The concomitant decrease in peroxiredoxin-2 (0.19-fold) is consistent with a reduced antioxidant capacity. Consistent with this hypothesis, soluble vitreous iron was lower in staphylomatous eyes, but not in highly myopic eyes without staphyloma, independently of total protein content and only weakly related to axial length; together with high vitreous ferritin and tissue iron deposits, this pattern suggests that iron is retained and stored in the staphylomatous retina rather than released into the vitreous. The lower vitreous levels of haemoglobin and α1-microglobulin in staphylomatous eyes are more difficult to interpret and may reflect differences in blood contamination at surgery rather than a disease mechanism.

In contrast, the decrease in neuron-derived proteins (brevican, chromogranin A, oligodendrocyte-myelin glycoprotein) and in interphotoreceptor retinoid-binding protein (RBP3) was already present in highly myopic eyes without staphyloma. RBP3 is secreted by photoreceptors and is required for their survival^17^, and its vitreous level falls in early diabetic retinopathy, when photoreceptor function is impaired^18^. Its decrease, like that of neuronal proteins, therefore probably reflects the reduction in photoreceptor density and retinal thickness that accompanies axial elongation^5^, rather than a feature specific to the staphyloma. Among glial markers, vimentin increased in the vitreous as in the tissue, whereas GFAP did not; as a cytoskeletal protein, GFAP is probably poorly released into the vitreous, so that its tissue increase is better assessed histologically.

Sirius red staining further showed that the scleral collagen lamellae were thinned and disorganized in highly myopic eyes, most markedly beneath the staphyloma. This agrees with the scleral thinning, loss of scleral tissue and weakening of its mechanical properties that accompany axial elongation and ultimately lead to posterior staphyloma^19^, and with the histomorphometric thinning of the staphylomatous sclera^6^. In the vitreous, the collagen module tended to increase, driven by procollagen C-endopeptidase enhancer 1 (PCPE-1) and collagen XVIII. PCPE-1 enhances the C-terminal maturation of fibrillar procollagens by BMP-1/tolloid-like proteinases and determines the ultrastructure of collagen fibrils in connective tissues^20^; its increase suggests an active, possibly aberrant, turnover of fibrillar collagen. Collagen XVIII is present in ocular basement membranes, including the inner limiting membrane, where it anchors vitreous collagen fibrils, and its loss causes high myopia, vitreoretinal degeneration and retinal detachment in Knobloch syndrome^21^. Because both proteins were already increased in highly myopic eyes without staphyloma, the extracellular matrix remodelling detected in the vitreous probably accompanies axial elongation, the staphyloma representing its most advanced stage. Our data do not establish whether vitreous PCPE-1 originates from the sclera, the choroid or retinal glial cells.

Our results only partly overlap with previous vitreous proteomic studies of pathologic myopia. Wen et al. reported higher levels of tubulin α1A and elongation factor 1α1, and lower levels of xylosyltransferase 1, versican and testican-2 in pathologic myopia^13^; we did not quantify the first three proteins, but versican tended to decrease in staphylomatous vitreous. Wei et al. described changes in antioxidant proteins^12^, which we did not reproduce. These studies grouped highly myopic eyes with various complications and did not separate eyes with and without staphyloma; differences in case definition, controls and quantification pipelines likely explain the limited overlap. Collagen XVIII and procollagen C-endopeptidase enhancer 1, increased in our myopic eyes, add to the evidence that extracellular matrix remodelling is reflected in the vitreous^13^.

This study has limitations. The proteomic series is small (4 controls, 3 highly myopic eyes without staphyloma and 7 with staphyloma), and no protein reached a 5% false discovery rate; the module analysis was defined a priori to limit multiple testing, but its results should be considered exploratory and need validation, for example by targeted assays of C1q, C4, factor D and ferritin in a larger cohort. Vitreous was obtained during vitrectomy for different indications, and the controls, operated for intraocular lens subluxation or epiretinal membrane, are not normal eyes; epiretinal membranes themselves involve glial proliferation. The absence of staphyloma was not documented with the same imaging in all highly myopic eyes, and in the iron series staphyloma status was assigned from clinical records, with only three eyes having a documented absence of staphyloma. Zero values were replaced by half the lowest observed abundance, which makes fold changes of proteins nearly absent from one group dependent on that floor. Finally, the histological and proteomic analyses were performed on different eyes and the histology is qualitative, so that the link between tissue and vitreous rests on convergence rather than on paired measurements.

In conclusion, the posterior staphyloma combines neuronal loss, glial scarring, microglial accumulation, iron overload and scleral collagen disorganization, and the vitreous of staphylomatous eyes carries a matching signature of classical complement activation and iron sequestration, distinct from the neuronal protein loss shared with high myopia. Complement and iron metabolism emerge as candidate pathways, and potential therapeutic targets, in the degeneration associated with posterior staphyloma. degeneration associated with posterior staphyloma.

## Methods

### Ethics statement

This study adhered to the tenets of the Declaration of Helsinki. Vitreous samples were collected at the Hospital Universitario Puerta de Hierro-Majadahonda (Spain) with the approval of **(ethics committee, reference number**), and all patients gave written informed consent. Human eyes used for histology were obtained from **(source**) with the approval of (**ethics committee or tissue bank, reference number)**.

### Patients and vitreous sampling

Undiluted vitreous was collected at the beginning of pars plana vitrectomy in patients operated for intraocular lens subluxation, epiretinal membrane, lamellar or full-thickness macular hole, vitreomacular traction or myopic traction maculopathy. High myopia was defined as an axial length of at least 26 mm or a refractive error of −6 dioptres or more. Posterior staphyloma was identified on (**imaging modality: optical coherence tomography, wide-field fundus imaging and/or magnetic resonance imaging**) and classified according to Curtin^3^. Eyes were assigned to three groups: non-myopic controls, highly myopic eyes without documented staphyloma (MS−) and highly myopic eyes with posterior staphyloma (MS+). Samples were stored at −80 °C until analysis. Patient characteristics are given in Table 1 and Supplementary Table S1.

### Histology and immunohistochemistry

[Number] highly myopic eyes with posterior staphyloma and [number] control eyes were fixed in [fixative], embedded in [paraffin], and serially sectioned at [thickness] µm along the temporal–nasal axis through the macula and the optic nerve head. Sections were stained with cresyl violet (Nissl), Perls’ staining for ferric iron, and picrosirius red for collagen. For immunofluorescence, sections were incubated with antibodies against Iba1 **([supplier, catalogue number, dilution]**), vimentin **([…])**, GFAP ([…]) and ferritin ([…]), followed by Alexa Fluor-conjugated secondary antibodies **([…])**; nuclei were counterstained with DAPI. Images were acquired with **XXXX** using identical settings for control and myopic sections.

### Proteomic sample preparation and LC-MS/MS

Proteomic analyses were performed by the PISSARO proteomics facility (Univ Rouen Normandie, France). Protein concentration was determined by the Bradford assay (bovine serum albumin standard, 0.1–1 mg/mL; absorbance at 595 nm) and checked on silver-stained 12.5% SDS-PAGE. For each sample, 12 µg of protein was concentrated in a 5% polyacrylamide stacking gel, stained with Coomassie blue, excised and digested overnight at 37 °C with trypsin (1 µg trypsin per 12 µg protein). Peptide concentration was determined by a peptide assay **[kit],** and 0.3 µg of peptides was injected onto a [**nano-LC system and column**] coupled to an Orbitrap Eclipse Tribrid mass spectrometer (Thermo Fisher Scientific). Data were acquired in data-dependent mode over a 150-min gradient with higher-energy collisional dissociation, Orbitrap detection of precursor (resolution 120,000, m/z 400–1800) and fragment ions **[authors to confirm with the facility]**.

### Protein identification and label-free quantification

Raw files were processed with Progenesis QI for Proteomics (Nonlinear Dynamics, Waters). Peptide ion maps were aligned to a reference run and normalized, normalization factors between 0.5 and 2 being considered acceptable since peptide amounts had been equalized before injection. Spectra were searched with [**search engine**] against the UniProtKB/Swiss-Prot Homo sapiens database, with trypsin specificity, **[number]** missed cleavages **and [fixed and variable modifications]**; identifications were filtered at a false discovery rate below 1%. Sample quality was assessed by intra-group correlation and alignment scores: one control with low intra-group reproducibility and two MS− samples with a low alignment score or a high normalization factor were excluded before analysis.

### Proteomic data analysis

Normalized protein abundances were exported from Progenesis QI. Proteins identified by a single peptide were excluded. Zero values were replaced by half of the lowest non-zero abundance of the same protein, and data were log2-transformed. Global structure was examined by principal component analysis of z-scored log2 abundances. Group differences were tested with an empirical-Bayes moderated t-test (limma approach), and p values were adjusted with the Benjamini–Hochberg procedure. Fold changes are ratios of geometric means. Welch’s t-test, the exact Mann–Whitney U test and bias-corrected Hedges’ g were computed as sensitivity analyses. Proteins with p < 0.05 and a fold change of at least 1.5 were considered regulated. Proteins detected in at least 5 of 7 samples of one group and in at most one sample of the other were additionally compared by Fisher’s exact test on detection frequency.

To relate vitreous changes to histological findings, nine protein modules were defined before testing from the known cellular origin or function of their members: classical complement, alternative and terminal complement, myeloid/lysosomal proteins, macroglial reactivity, neuronal/synaptic proteins, photoreceptor and interphotoreceptor matrix proteins, collagens and procollagen processing, iron transport and storage, and antioxidant enzymes (Supplementary Data 1). Every member identified by at least two peptides was included regardless of its individual result. For each sample, a module score was computed as the mean of the per-protein z-scores. Differences in module scores were tested by exact permutation of group labels (all possible relabelings, two-sided), with Benjamini–Hochberg correction across modules, and robustness was assessed by leave-one-sample-out recomputation. In a secondary analysis, the MS− eyes, quantified in the same run with identical normalization, were added and a three-group moderated t-test with pooled variance was applied. Associations between proteins and module scores were assessed with Spearman’s correlation. Analyses were performed in Python 3.13 with SciPy and statsmodels.

### Vitreous iron and protein assays

In an independent series of vitreous samples assayed in a single batch, soluble iron was measured by **[method, kit, supplier]** and total protein by **[method].** Iron was also expressed relative to total protein (ng of iron per mg of protein). Groups were compared with the Mann–Whitney U test and the Kruskal–Wallis test, and associations with axial length with Spearman’s correlation. All tests were two-sided and p < 0.05 was considered significant.

## Data availability

The mass spectrometry proteomics data have been deposited to the ProteomeXchange Consortium via the PRIDE partner repository with the dataset identifier **[PXD000000].** Processed protein abundances and statistical results are provided in Supplementary Data 1, and patient characteristics in Supplementary Table S1.

---

**Supplementary Figure S1.**
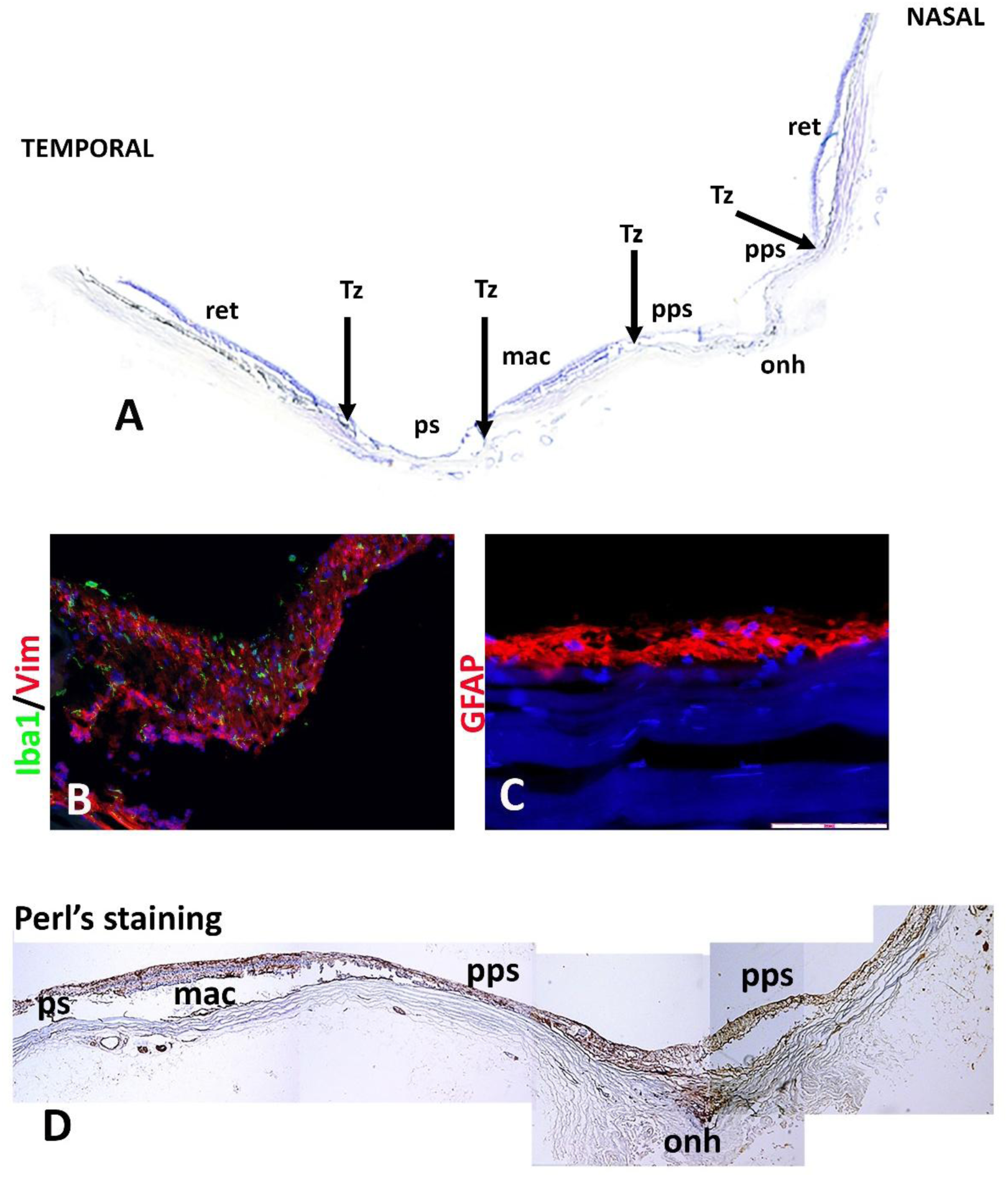
Topography of the posterior staphyloma. (**a**) Nissl-stained section along the temporal–nasal axis of a highly myopic eye, showing the temporal retina (ret), transition zones (Tz), posterior staphyloma (ps), macula (mac), peripapillary staphyloma (pps) and optic nerve head (onh). (**b**) Iba1 (green) and vimentin (red) immunofluorescence at a transition zone. (**c**) GFAP immunofluorescence (red) in the staphyloma, nuclei in blue. (**d**) Perls staining from the posterior staphyloma to the optic nerve head (composite of adjacent fields). Scale bars: [authors to add].

**Supplementary Figure S2.**
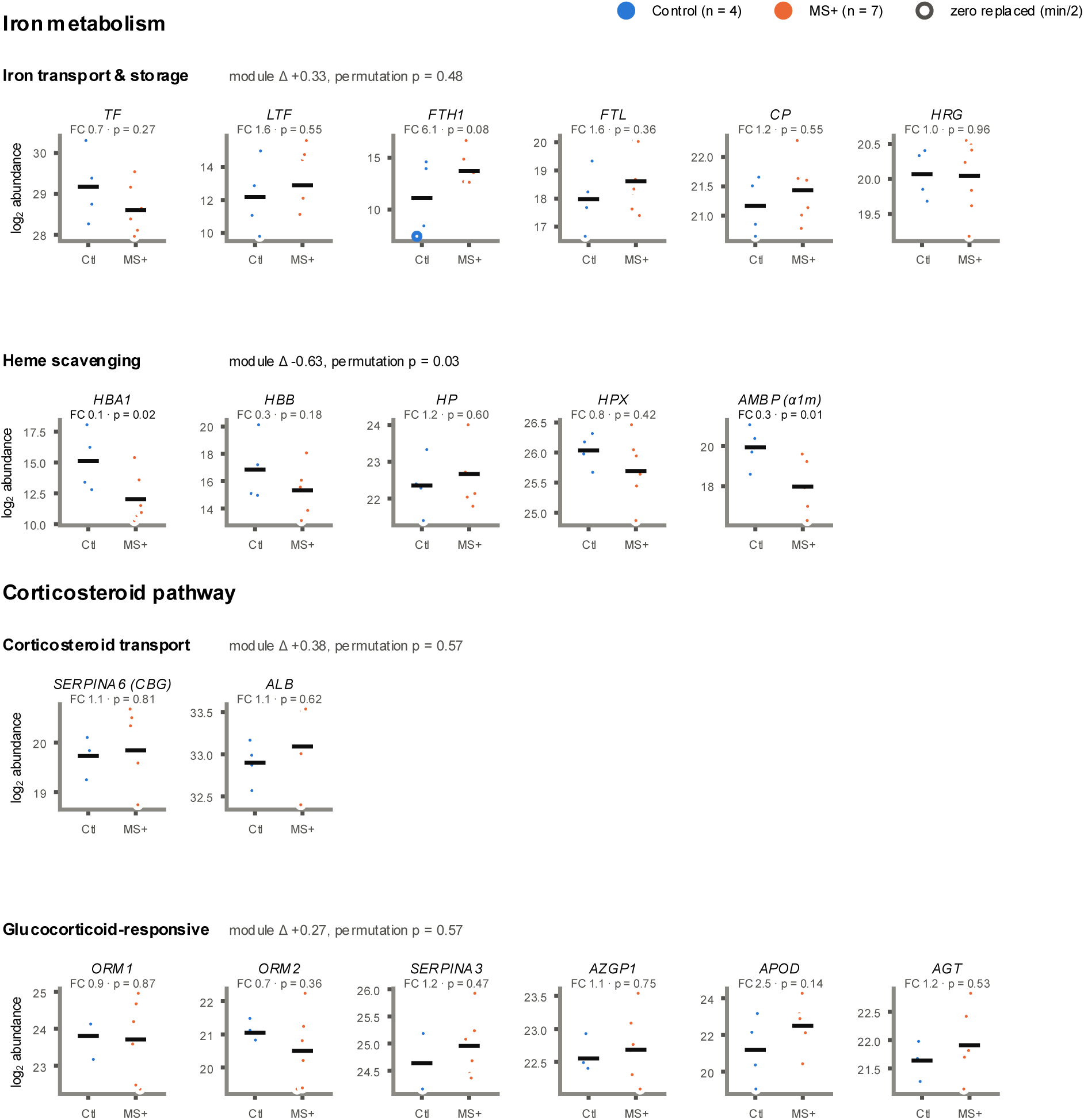
Iron metabolism and corticosteroid pathway in the vitreous proteome. Iron transport and storage, haem scavenging, corticosteroid transport and glucocorticoid-responsive proteins in controls (n = 4) and MS+ eyes (n = 7). Points, individual eyes; bars, means. Fold change and moderated t-test p value above each panel; module score difference and exact permutation p value beside each row title.

**Supplementary Figure S3.**
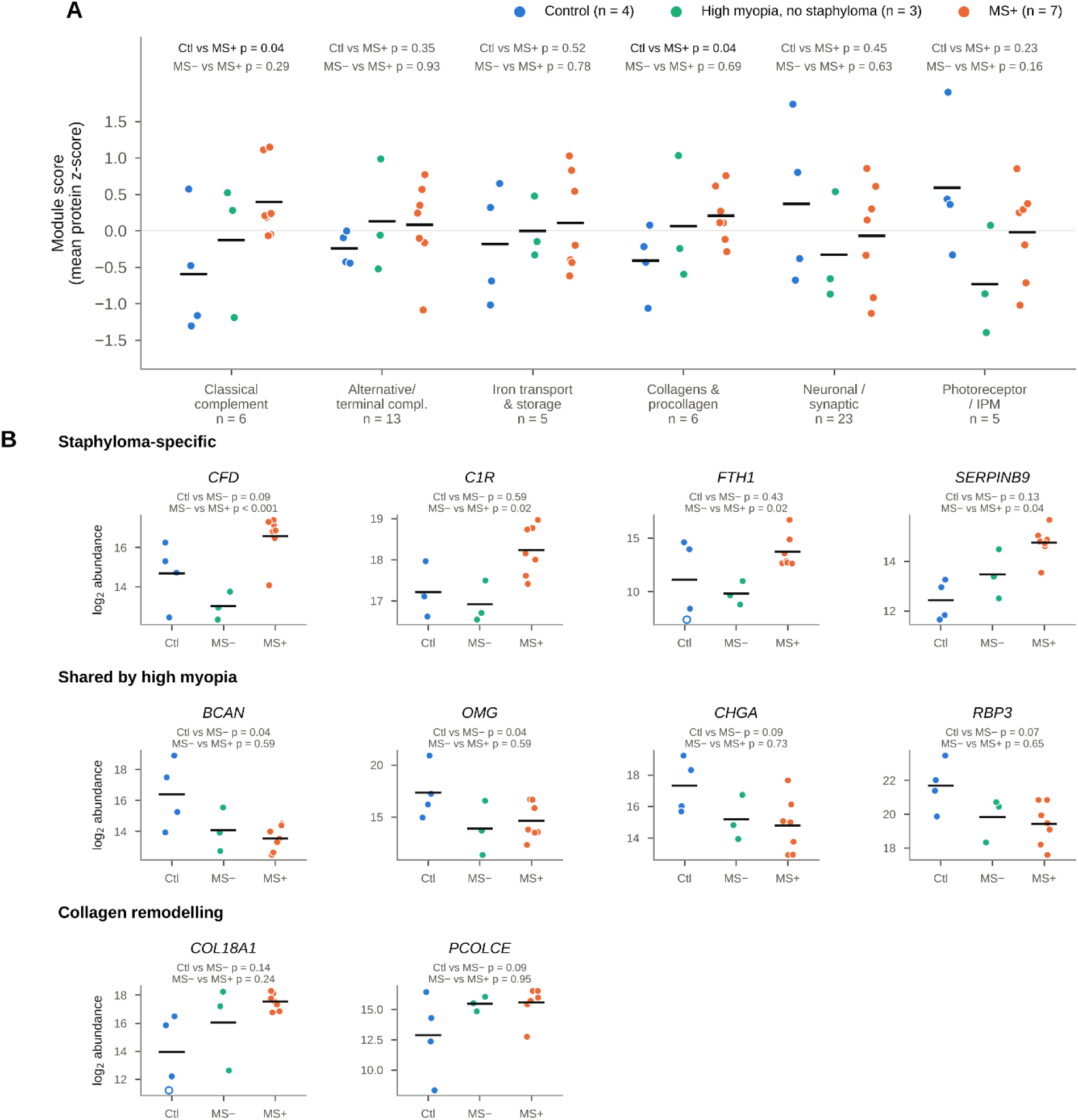
Vitreous changes specific to the staphyloma versus shared with high myopia. (**a**) Module scores in controls (blue, n = 4), highly myopic eyes without documented staphyloma (MS−, green, n = 3) and MS+ eyes (orange, n = 7), with exact permutation p values for control versus MS+ and MS− versus MS+. Scores are z-scored across the 14 eyes and therefore differ slightly from Fig. 3a. (**b**) Proteins higher in MS+ than in MS− vitreous, proteins already decreased in MS− vitreous, and collagen-processing proteins. Moderated t-test p values from a three-group model; open circles, replaced zero values.

## Supplementary Information

| File | Content |
|---|---|
| Supplementary Table S1 (Excel) | Patient characteristics. Sheet A: the 17 eyes submitted to proteomics, with inclusion status, sex, age, axial length, staphyloma and indication. Sheet B: the 31 eyes of the vitreous iron series. |
| Supplementary Data 1 (Excel) | Proteomic acquisitions and results: all identified proteins with normalized abundances for the 14 analysed eyes; control versus MS+ analysis; three-group analysis; module definitions, tests and per-eye scores; analysis methods. |
| Supplementary Figures S1–S3 | S1, topography of the posterior staphyloma (Nissl, Iba1/vimentin, GFAP, Perls); S2, iron and corticosteroid pathways in the vitreous proteome; S3, staphyloma-specific versus high-myopia changes. |

